# PrISM: Precision for Integrative Structural Models

**DOI:** 10.1101/2021.06.22.449385

**Authors:** Varun Ullanat, Nikhil Kasukurthi, Shruthi Viswanath

## Abstract

**Motivation:** A single precision value is currently reported for an integrative model. However, precision may vary for different regions of an integrative model owing to varying amounts of input information.

**Results:** We develop PrISM (Precision for Integrative Structural Models), to efficiently identify high and low-precision regions for integrative models.

**Availability:** PrISM is written in Python and available under the GNU General Public License v3.0 at https://github.com/isblab/prism; benchmark data used in this paper is available at doi:10.5281/zenodo.6241200.

**Supplementary information:** Supplementary data are available at *Bioinformatics* online.

## 1 Introduction

Integrative modeling has emerged as the method of choice for determining the structures of macromolecular assemblies which are challenging to characterize using a single experimental method (Alber *et al*., 2007; Russel *et al*., 2012; Ward *et al*., 2013; Webb *et al*., 2018; Rout and Sali, 2019; Saltzberg *et al*., 2021). Several assemblies have been determined by this approach, yielding insights on transcription (Robinson *et al*., 2015), gene regulation and DNA repair (Luo *et al*., 2015; Arvindekar *et al*., 2021), intra-cellular transport (Kim *et al*., 2018; Ganesan *et al*., 2020), cell cycle progression (Viswanath, Bonomi, *et al*., 2017; Pasani and Viswanath, 2021), immune response and metabolism (Lasker *et al*., 2012; Gutierrez *et al*., 2020). Integrative modeling often relies on sparse, noisy, and ambiguous data from heterogenous samples (Schneidman-Duhovny *et al*., 2014). Usually, more than one model (structure) that satisfies the data. Therefore, an important attribute of an integrative model is its precision, defined as the variability among the models that satisfy the input data. The precision defines the uncertainty of the structure and is a lower bound on its accuracy. Importantly, downstream applications of the structure are limited by its precision. For example, a protein model of 20 Å precision cannot be used to accurately identify binding sites for drug molecules. Precision aids in making informed choices for future modeling, including the representation, degrees of freedom, and the amount of sampling (Viswanath, Chemmama, *et al*., 2017; Pasani and Viswanath, 2021).

Currently, a single precision is reported for the integrative model. However, there can be varying amounts of input information for different regions in the model, resulting in different precisions for different regions (Viswanath and Sali, 2018). It would be useful to identify regions of high and low-precision in the model. For instance, low-precision regions can suggest where the next set of experimental data would be most impactful. High-precision regions can be used for further analysis such as identifying binding interfaces, rationalizing known mutations, and suggesting new mutations.

Several methods have been proposed for detecting substructure similarities and determining flexible/rigid regions (Wriggers and Schulten, 1997; Kedem *et al*., 1999; Jacobs *et al*., 2001; Pfleger *et al*., 2013; Martínez, 2015; Cazals and Tetley, 2019). However, they are not directly applicable to integrative models of macromolecular assemblies. First, they rely on the input being a set of atomic structures with known secondary structure. In contrast, integrative models are encoded by a more complex representation (ensemble of multi-scale, multi-state, time-ordered models), and can comprise of regions with unknown structure (Viswanath and Sali, 2018; Sali *et al*., 2015; Vallat *et al*., 2018). Second, these methods identify rigid substructures without quantifying precision for all parts of the structure. Finally, these methods analyze structures with a small number of proteins and have not been demonstrated to be scalable for large ensembles of macromolecular assemblies.

Validation of integrative models, including assessment of model precision, is an open research challenge and timely due to the new PDB archive for integrative structures (Sali *et al*., 2015; Vallat *et al*., 2018; Berman *et al*., 2019; Vallat *et al*., 2019, 2021) (http://pdb-dev.wwpdb.org). Here, we demonstrate PrISM, a method to visualize high and low-precision regions of an integrative model. Methods like PrISM are expected to improve the utility of deposited integrative structures.

## 2 Methods

### 2.1 PrISM Inputs and Outputs

The input is a set of structurally superposed integrative models (Fig. 1). Commonly, these models are encoded by a multi-scale representation, although PrISM also supports integrative models in the atomic representation in PDB format (Viswanath and Sali, 2018; Sali *et al*., 2015; Vallat *et al*., 2018). In the multi-scale representation, each protein is represented by a sequence of spherical beads; each bead corresponds to a number of contiguous residues along the protein sequence. Coarse-grained bead representations are necessary since large assemblies cannot be efficiently and exhaustively sampled in atomic detail (Viswanath and Sali, 2018; Rout and Sali, 2019; Saltzberg *et al*., 2021). Regions with atomic structure are represented at higher-resolution (e.g., one residue per bead); other regions are usually further coarse-grained (e.g., thirty residues per bead). The most common input would be the models from the most populated cluster from integrative modeling analysis (Viswanath, Chemmama, *et al*., 2017; Saltzberg *et al*., 2019, 2021). Additional optional user inputs include the voxel size for bead grids and the number of high and low-precision classes.

**Fig. 1.**
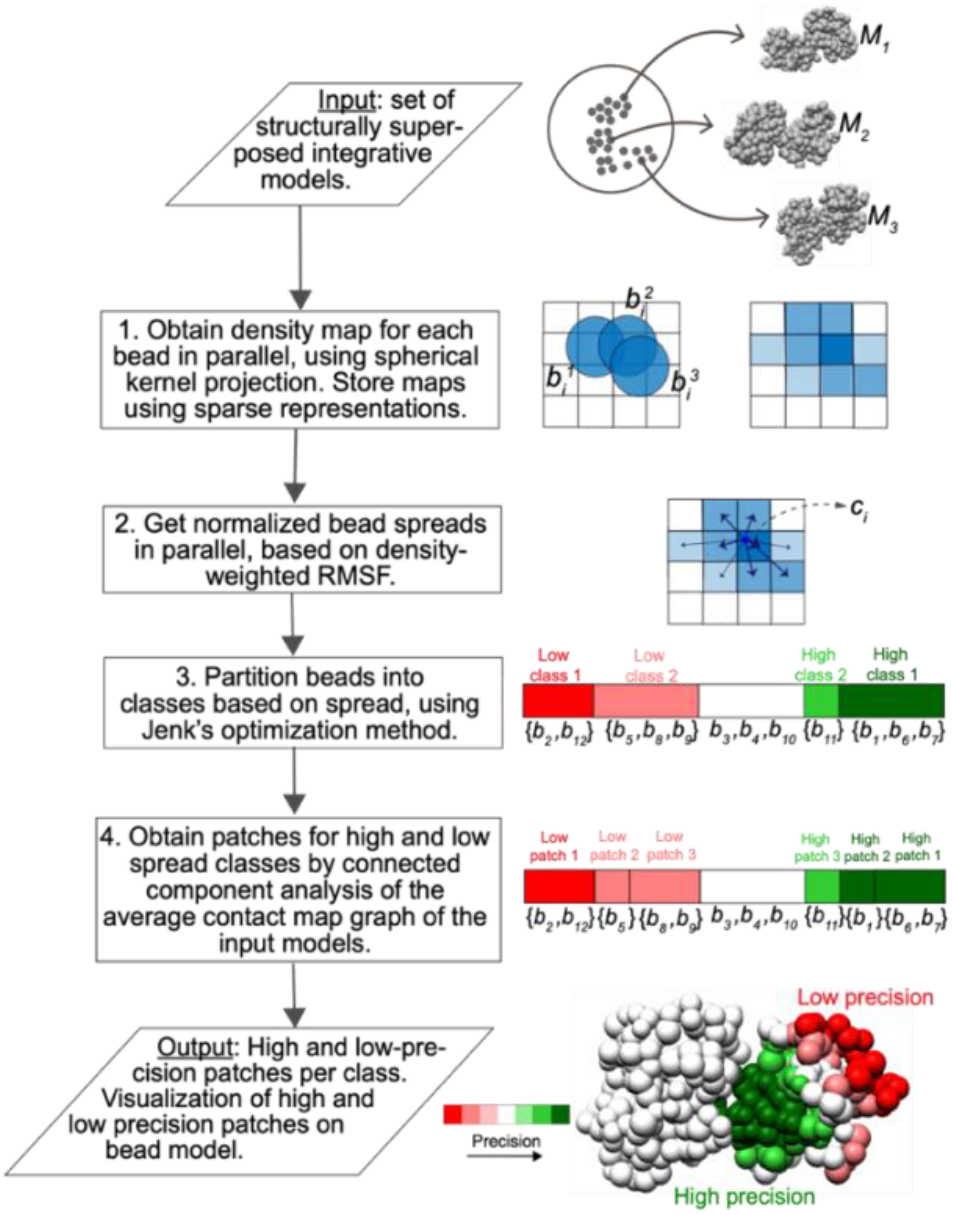
Schematic of PrISM. The input is a set of structurally superposed integrative models. Models of three protein-protein complexes are depicted here as *M*_1_, *M*_2_ and *M*_3_. First, a density map is obtained for each bead. Three beads corresponding to bead *i* from the three models, *M*_1_, *M*_2_ and *M*_3_, are projected onto the grid. The obtained density map has blue-colored squares; the color intensity corresponds to the density of the square. Next, the normalized bead spread is computed from the density map as the deviation of densities around the center of density *c*_*i*_. Subsequently, the Jenks method is used to classify beads into high and low-precision classes. In the example shown, there are two low and two high-precision classes. These classes are further partitioned into patches. The output is a set of high and low-precision patches per class. It is visualized on a representative model as a bi-polar colormap, with shades of green (red) corresponding to high-precision (low-precision) patches.

The outputs from PrISM are regions (‘patches’) of high and low-precision. They are visualized as a bi-polar color map overlaid on a representative model, with high (low)-precision patches in shades of green (red).

### 2.2 PrISM Algorithm

The algorithm is described here. Alternate design choices are also discussed (Supplementary Methods).

#### 2.2.1 Obtaining bead-wise density maps

A coarse-grained bead is the smallest primitive, *i*.*e*., unit of representation, of an integrative model. We first compute a density map for each bead. A density map is a projection or rasterization of the beads onto a 3D grid, storing a density value for each grid element (voxel). We use a spherical kernel projection since it explicitly considers the bead mass and radius. The contribution to density 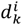 to voxel *k*, centered at *v*_*k*_, in a grid with voxel spacing *V*, from bead *i* of *m*odel *M*_*j*_, with centre coordinates 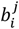, mass *m*_*i*_, and radius *r*_*i*_ is given by: 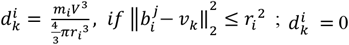 otherwise.

The densities at each voxel are subsequently normalized by the number of input models to obtain the average density at a voxel.

Since the density map for each bead can be independently computed, this step is trivially parallelized. The density map provides a uniform representation for comparing beads of different sizes.

#### 2.2.2 Computing bead spread

We define the bead spread, a measure of bead precision, as the density-weighted RMSF from the bead center of density. That is, the density center *c*_*i*_ for bead *i* is: 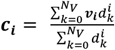 where *N*_*v*_ is the number of voxels in the grid. Bead spread *s* is computed by: 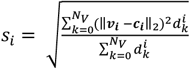. This step is also parallelized. The bead spreads are then normalized to 0 to 1 using min-max scaling.

#### 2.2.2 Classifying beads by spread

Next, we use the Jenks Natural Breaks algorithm to classify beads into high and low-precision classes, given the required number of high- and low-precision classes (Jenks, 1967). This algorithm produces the classification that optimizes a goodness-of-variance measure, similar to k-means clustering. It is used in thematic mapping for clustering one-dimensional data (Jenks, 1967).

### 2.2. Obtaining patches

Next, we detect beads with concerted localization by identifying correlations in the positions and precisions of beads. A patch is a set of beads in the same precision class, computed by the Natural Breaks algorithm above, which are also proximal to each other in the set of models. A pair of beads is proximal if the average distance between their surfaces, across the input set of models, is less than 10 Å. Patches represent a further grouping of beads in each class. We obtain patches efficiently by deriving connected components of the graph representing the average contact map of the input set of models. A naïve implementation of the above algorithm is prohibitively expensive in terms of runtime and memory even for small complexes, necessitating several software enhancements (Supplementary Methods).

## 3 Evaluation and Usage

PrISM is benchmarked on twelve systems and shown to be fast (Supplementary Results, Table S1) (Viswanath, Chemmama, *et al*., 2017; Saltzberg *et al*., 2019, 2021; Viswanath and Sali, 2018; Brilot *et al*.; Luo *et al*., 2015; Arvindekar *et al*., 2021). The annotated precision is shown to be consistent with root mean-square fluctuation (RMSF) and localization density maps, providing more fine-grained information than the latter in some cases (Supplementary Results, Table S2, Fig. S1-S3). We recommended parameters for PrISM (Supplementary Results, Table S3, Fig. S4-S5). Finally, we explain how PrISM output can be used to distinguish between conformational heterogeneity, *i*.*e*., multiple states, and lack of data (Supplementary Results).

Detailed usage is at https://github.com/isblab/prism.

## 4 Conclusion

PrISM is an efficient method for annotating precision for integrative models of large assemblies. A limitation is that it is applicable to structurally superposed atomic models (generated by any integrative modeling software) and integrative models generated by the Integrative Modeling Platform (IMP, https://integrativemodeling.org). In contrast to atomic structural models, models from IMP are multi-scale, coarse-grained at multiple levels by spherical beads. In future, the approach could be extended to other model ensembles of coarse-grained models. Methods such as PrISM are expected to improve the utility of deposited integrative structures in the PDB (http://pdb-dev.wwpdb.org) (Sali *et al*., 2015, 20; Vallat *et al*., 2018, 2019, 2021).

## Acknowledgements

We thank Shreyas Arvindekar, Satwik Pasani, Aditi Pathak, Kartik Majila, and Praveen Roy DS for feedback on the manuscript and testing out early versions of the method.

