## Supplementary Material for "PrISM: Precision for Integrative Structural Models"

Short title: Precision for integrative models

*Keywords*: model precision, integrative structure modeling, validation of integrative models, density map, 3D grid, spatial clustering, thematic mapping.

### Supplementary Methods

#### 1. Algorithm enhancements for runtime and memory usage efficiency

A naïve implementation of PrISM is prohibitively expensive in terms of runtime and memory even for small complexes. So we implemented the following enhancements. First, comparing densities across beads requires that all beads be projected onto the same 3D grid (*global* grid), whose dimensions (bounding box) correspond to the maximum span of the coordinates in each dimension (*X*, *Y* and *Z*) across all beads and all input models. To reduce the runtime, we first project a bead onto a bead-wise grid (*local* grid) and then map the projection onto the global grid.

Second, we make the density calculation less memory-intensive. Projecting a given bead onto a 3D grid requires computing the distance matrix between the coordinates of the given bead in all input models and all the voxels of the grid. This operation becomes memory intensive due to an exponential increase size of the distance matrix when the number of voxels is large. The latter can occur when the local grid dimensions are large and/or when the voxel size is small. To reduce the memory requirements, we divide set of possible voxels into batches and compute the distance matrix separately for each batch while preserving voxel identity. A pitfall is that this increases the computation time slightly.

Third, we use a sparse representation for the bead densities, storing only voxels associated with non-zero density, thus decreasing the memory required and runtime for subsequent calculations.

Fourth, we also provide options to set the voxel size of the density maps and the percentage of input models to use. Larger voxels correspond to lower runtime and memory requirement. The runtime can be decreased by randomly sampling a subset of input models, instead of all models.

Finally, one can reduce the number of beads to be annotated by selecting beads at the coarsest resolution of a multi-scale model and not selecting beads that are fixed during sampling.

#### 2. Alternate design choices

**Alternate kernels** Other kernels such as Gaussian kernels could be used to project beads to density maps. However, these kernels treat beads as point masses and do not consider the bead radii explicitly.

**RMSF instead of bead spread** Similarly, RMSF could be considered instead of bead spread. However it only considers bead coordinates and does not account for the mass or size of the bead. Moreover, RMSF is a bead-wise measure and is not sufficient by itself. Additional computation would be required to identify correlations between beads in order to group them to obtain regions of high and low-precision.

**Alternate methods to classify beads** Other methods could be used to group beads based on spread. In particular, we also considered grouping beads based on the inter-quartile range (IQR) of bead spread. Beads with spreads at 25 percentile and lower were marked as high precision and those with spreads at 75 percentile and higher were marked as low-precision. However, due to the hard cut-offs at 25 and 75 percentiles, slight differences in the bead spread resulted in adjacent beads in rigid bodies being classified differently. The Jenks method, on the other hand, resulted in a more uniform classification for beads in rigid bodies.

#### 3. Datasets

A total of twelve systems were used in the study. Five systems were binary protein-protein complexes that formed the benchmark in a previous study on protocols for analysing integrative models: complex between trypsin and its inhibitor (PDB 1AVX), RAN and RCC1 (PDB 1I2M), colicin and its inhibitor (PDB 7CEI), spliceosomal protein and its inhibitor (PDB 1SYX), and subunits of DNA polymerase III (PDB 2IDO) [1], [2]. The other systems included the complex between actin, gelsolin and tropomyosin [3], yeast γ-tubulin small complex bound to Spc110 [4], the transcription and DNA repair factor TFIIH [5], RNA polymerase II [6], and three sub-complexes of the nucleosome remodeling and decetylase complex [7]. These datasets are in public repositories on https://integrativemodeling.org. For each system, the input was the set of models from the most populated cluster from IMP analysis.

### Supplementary Results

#### 1. Runtime

The runtime of PrISM was benchmarked; it takes a few minutes to run PrISM on a modern workstation (Table S1). The runtime increases linearly with the fraction of input models used and decreases with voxel size (Fig. S3).

#### 2. Comparison to RMSF

Next, we check for consistency of the bead spread with the bead-wise RMSF (root mean-square fluctuation), using the Spearman rank correlation [8] [1]. The RMSF only considers the deviation of bead coordinates, while the bead spread additionally accounts for the bead mass and radius. Nevertheless, a positive correlation between the two exists for all the studied systems, with a strong correlation for most systems (Table S2, Fig. S1).

#### 3. Comparison to localization density maps

Next, the annotated high and low-precision patches were also qualitatively compared with the localization probability density maps of the cluster [1], [7], [9], [10] (Fig. S2). Localization density maps specify the probability of any volume element being occupied by a given bead in superposed models. In the first example, a set of integrative models is obtained by docking a monomer of Spc110-N terminus to the fixed γ-tubulin small complex (γ-TuSC) using chemical crosslinks, cryo-EM, and stereochemistry information [10]. The precision-colored model from PriSM and the localization density map are compared for Spc110-N terminus (Fig. S2A-S2B).

The Spc110$^{164-203}$ helix is kept fixed during simulation. Therefore its localization is precise in the density maps (Fig. S2A). This is consistent with the PrISM output where the helix is annotated as a high-precision region (Fig. S2B). On the other hand, Spc110$^{1-163}$ is predicted to be disordered and represented by flexible beads (Fig. S2A). The corresponding region in the precision-colored model shows beads of varied precision, including some at high-precision. This indicates that at least some regions in the disordered N-terminus of Spc110 can be precisely localized on γ-TuSC based on the input information. Interestingly, these precise regions correspond to the conserved centrosomin domain of Spc110 visualized in newer cryo-EM maps [4]. This information, while not available from the localization density maps, is available from the PrISM output as it is a more fine-grained visualization of the ensemble.

The subsequent examples comprise of models of sub-complexes of the nucleosome remodeling and deacetylase (NuRD) complex based on negative-stain EM, chemical crosslinks, and stereochemistry information [7]. In MHR, the densities of R1-RBBP4 are spread out, while those of R2-RBBP4s are precisely localized, consistent with the PrISM output (Fig. S2C-S2D). Finally, in the NuDe complex, the densities of the RBBP4 on the right are spread out, while those of MTA1-HDAC1 dimer and MBD3 are precisely localized, consistent with the PrISM output (Fig. S2E-S2F). In summary, the precision annotation from PrISM is broadly consistent with localization density maps, while providing more fine-grained information than the latter, for some systems.

#### 4. Recommended parameters

Finally, we recommend values of voxel size and number of Jenks classes, based on the examined benchmark (Table S3, Fig. S4-S5).

**Voxel size** On the examined complexes, bead spreads computed using a voxel size of 4 Å are very similar to those obtained using a smaller voxel size of 2 Å (Table S3, Fig. S4). Therefore, it is recommended to run PrISM with a voxel size of 4 Å.

**Number of classes** We recommend using two Jenks classes initially, as a balance between the discriminative power and ease of interpretability of the model. We compare the PrISM output with one, two, and three classes on the NuDe complex (Fig. S5).

#### 5. Distinguishing between multiple states and lack of data

It is possible to distinguish between multiple states and low resolution (or lack of data) using the PrISM output and an additional consideration, i.e., the fit to input data. If different models (conformations) satisfy different subsets of input data, one can conclude that there are multiple states. For example, models A and B in the input set of models may each satisfy 30% and 40% of the input chemical crosslinks respectively; in this situation, the system may contain a mixture of the two states. On the other hand, if different models satisfy all the input data, it could be a case of low-resolution or lack of data. For example, models A and B are different conformations, but both fit into the same low-resolution EM map.

#### Supplementary Figures

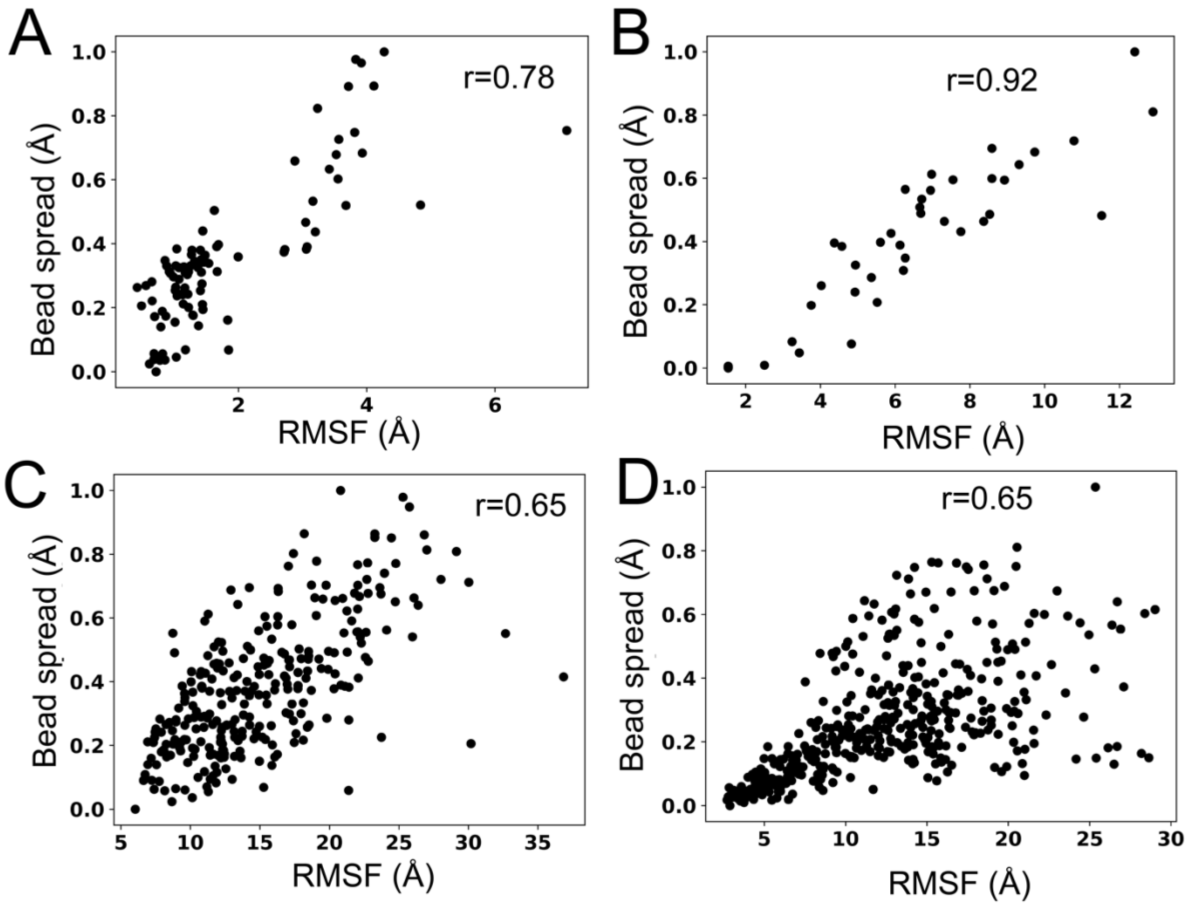


**Fig. S1 Correlation of bead spread with RMSF.** Scatter plot showing the Spearman rank correlation [8] between the RMSF and bead spread for all beads in (A) Actin, (B) Spc110-γTuSC, (C) MHR, (D) NuDe complexes.

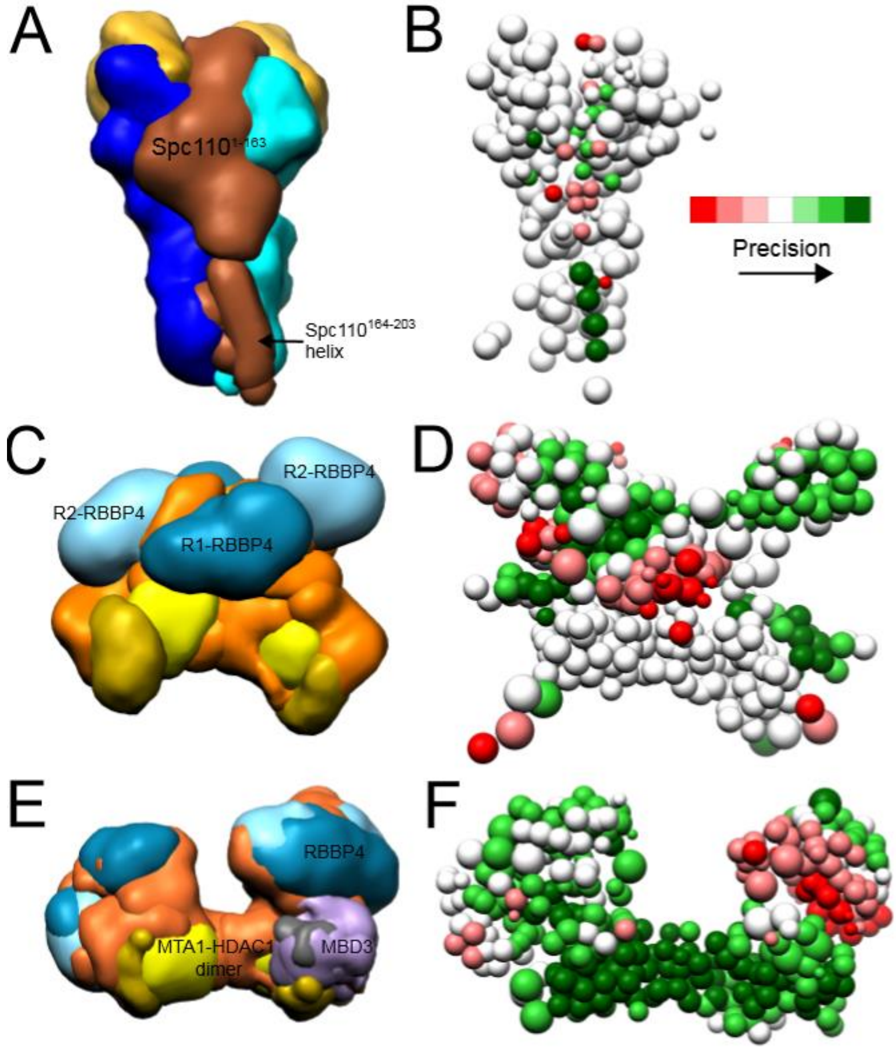


**Fig. S2 Comparison of PrISM output to localization density maps.** Localization density maps of the input set of models (A,C,E) and high and low-precision patches mapped on the representative bead models (B,D,F) for three systems are shown: the Spc110- γTuSC complex (A,B) (Brilot et al 2021), MHR (C,D), and NuDe (E,F) complexes [7]. All density maps were contoured at 10% of their respective maximum values.

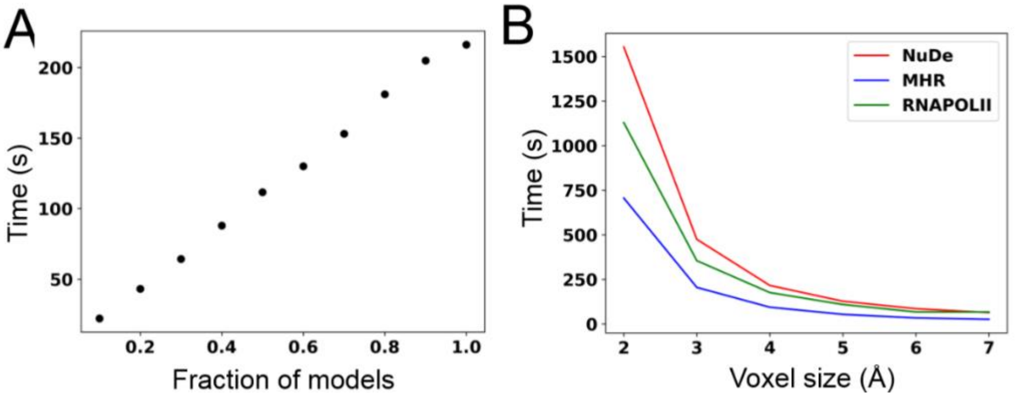


**Fig. S3 Runtime or PrISM.** The runtime of PrISM was visualized against (A) the fraction of input models for the NuDe complex, and (B) voxel size for NuDe, MHR and RNAPOLII complexes. The PrISM parameters used were: 4 Å voxel size and two Jenks classes. The benchmarking was performed on 16 cores of a second generation dual Intel Xeon Silver processer 4208 with 2.10 GHz clock speed.

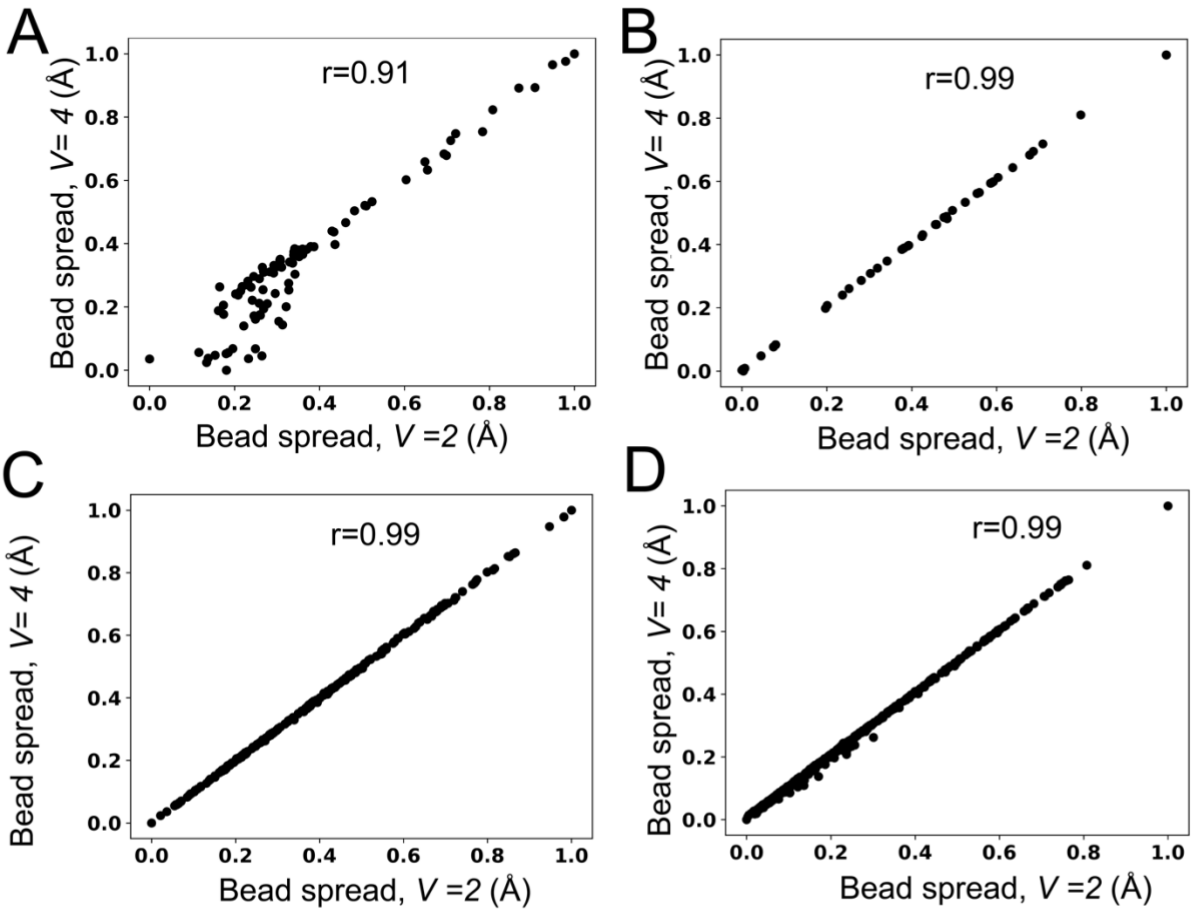


**Fig. S4 Correlation between bead spreads at voxel sizes 4 Å and 2 Å.** Scatter plot showing the Spearman rank correlation [8] between the bead spreads at voxel sizes 4 Å and 2 Å for all beads in (A) Actin, (B) Spc110- γTuSC, (C) MHR, (D) NuDe complexes.

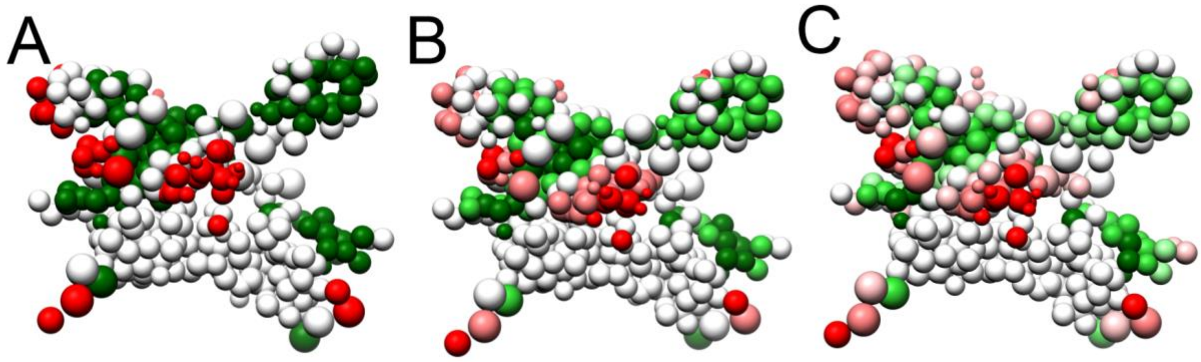


**Fig. S5 PrISM output for different numbers of Jenks classes.** The precision-colored bead model for the MHR complex was visualized for (A) 1, (B) 2, and (C) 3 Jenks classes. A voxel size of 4 Å was used throughout.

### Supplementary Tables

| Complex | Number of Beads | Number of Models | Runtime (seconds) +/- standard deviation |
|---|---|---|---|
| **ACTIN** | 92 | 1021 | 1.63±0.07 |
| **GTUSC** | 41 | 1621 | 1.36±0.06 |
| **MHM** | 222 | 13227 | 78.10±1.29 |
| **MHR** | 282 | 12913 | 91.19±1.19 |
| **NuDe** | 419 | 21625 | 219.54±3.05 |
| **RNAPOLII** | 453 | 11866 | 169.55±3.05 |
| **TFIIH** | 160 | 3203 | 25.05±0.27 |
| **1AVX** | 169 | 2897 | 2.13±0.04 |
| **1I2M** | 401 | 21612 | 42.80±0.42 |
| **1SYX** | 62 | 7241 | 1.69±0.04 |
| **2IDO** | 66 | 10462 | 2.48±0.04 |
| **7CEI** | 131 | 5921 | 2.82±0.04 |

**Table S1. Runtime of PrISM.** The average runtime of PrISM is shown against the number of beads in the system and the number of models in the ensemble over 10 repeated runs. The PrISM parameters used were: 4 Å voxel size and two Jenks classes. The benchmarking was performed on 16 cores of a second-generation dual Intel Xeon Silver processer 4208 with 2.10 GHz clock speed.

| Complex | Correlation |
|---|---|
| **ACTIN** | 0.78 |
| **GTUSC** | 0.91 |
| **MHM** | 0.88 |
| **MHR** | 0.65 |
| **NuDe** | 0.65 |
| **RNAPOLII** | 0.46 |
| **TFIIH** | 0.59 |
| **1AVX** | 0.92 |
| **1I2M** | 0.71 |
| **1SYX** | 0.51 |
| **2IDO** | 0.88 |
| **7CEI** | 0.86 |

**Table S2. Comparison of bead spread with RMSF.** The Spearman rank correlation [8] between the bead spread and the RMSF is shown for all the complexes. The PrISM parameters used were: 4 Å voxel size and two Jenks classes.

| Complex | Correlation |
|---|---|
| **ACTIN** | 0.91 |
| **GTUSC** | 0.99 |
| **MHM** | 0.99 |
| **MHR** | 0.99 |
| **NuDe** | 0.99 |
| **RNAPOLII** | 0.41 |
| **TFIIH** | 0.99 |
| **1AVX** | 0.92 |
| **1I2M** | 0.99 |
| **1SYX** | 0.99 |
| **2IDO** | 0.99 |
| **7CEI** | 0.99 |

**Table S3. Correlation between bead spreads at voxel sizes 4 Å and 2 Å.** The Spearman rank correlation [8] between the bead spread at voxel size 4 Å and 2 Å for all complexes are shown.

[10] A. F. Brilot *et al.*, “CM1-driven assembly and activation of yeast γ-tubulin small complex underlies microtubule nucleation,” *eLife*, vol. 10, p. e65168, doi: 10.7554/eLife.65168.
